# The extracellular RNA pool within *Zea mays* apoplast: composition and differential expression during *Ustilago maydis* infection

**DOI:** 10.1101/2022.06.03.494492

**Authors:** Dibya Mukherjee, Nagendra Pratap Singh, Anisha Roy, Rituparna Mondal, Udita Acharya, Debasis Chattopadhyay, Anupama Ghosh

## Abstract

The existence of an extracellular pool of RNA (exRNA) has been documented in both animal and plant cells in a number of instances. These exRNA species play important role in host response against different environmental stimuli. The mechanism of their function however remains largely unknown. In this study we report the composition of the exRNA pool within the leaf apoplast of *Z. mays* under normal growth condition. We could detect RNA transcripts originating from both the genic as well as the intergenic regions of the nuclear, mitochondrial and chloroplast genomes of maize in our exRNA sequencing data. Our data showed increased abundance of about 75% of the exRNA transcripts during infection with a basidiomycete smut fungi, *Ustilago maydis*. Functional classification of the differentially abundant exRNA transcripts within *U. maydis* SG200 WT infected maize apoplast with respect to uninfected apoplast revealed significant enrichment of the exRNA transcripts corresponding to the ribosome biogenesis pathway. Data related to the effect of two extracellular T2 type ribonucleases, Nuc1 and Nuc2 from *U. maydis* on the composition of exRNA pool of maize is also presented.

---

Plant apoplast remains a very important compartment involved in the response of plants toward environmental stimuli ^1^. It is a hub of defence responses upon exposure of a host plant to a stressful environmental condition and inhabits several defence proteins ^2^. In the recent past, however, RNAs as one of the major constituents of apoplasts have also been evidenced in some plants like Arabidopsis. These apoplastic RNAs are either confined within extracellular vesicles (EV) or are free-living within the apoplastic fluids. The extracellular pool of RNA (exRNA) within the Arabidopsis EVs is enriched in small RNAs including miRNAs, siRNAs, and a class of 10-17 nucleotides long RNA called tiny RNA species ^3^. A major population of the free-living exRNAs however comprises long non-coding RNAs including circular RNAs in Arabidopsis ^4^. Although not much is yet known about the biological function of this ex RNA pool in plants, some role in host defence against pathogen invasion has been anticipated. Silencing of pathogen genes through cross-kingdom gene silencing using EV enclosed small RNAs is one such example ^5^. However, the biological function of the free apoplastic RNAs both coding and non-coding remains elusive. One of the major concerns includes the stability of the free RNAs within the apoplastic space. Studies carried out in this direction revealed that various RNA binding proteins are involved in the stabilization of the free-living exRNAs that are not confined within protective vesicles ^4^. RNA binding proteins, therefore, are found to be extremely important for the sustenance of the free exRNAs within the apoplast. In a recent study, we have demonstrated the presence of a pool of RNA within the apoplast of maize ^6^. These exRNA species in maize exhibit a completely different gel profile compared to the total RNA from maize leaves when run on an 8% urea-polyacrylamide gel thereby indicating a separate composition than total transcriptome (Figure S1). This also points toward an as yet unknown mechanism through which a subpopulation of total RNA finds its path to the apoplast and is included in the exRNA pool. The composition of this exRNA pool from maize however remains unknown till date. In this study, we have now sequenced the apoplastic RNA pool of maize leaves under normal growth conditions and assessed the effects on its composition during infection with *Ustilago maydis*. However, our RNA isolation method from maize apoplastic fluid does not include a separate EV purification step. Due to this, it is not possible to classify the identified exRNA transcripts from maize apoplast into either EV or EV-free environment in this study. Our data revealed a total of 9599 transcript assemblies from both genic as well as intergenic regions from the maize genome within the apoplast under normal growth conditions. The transcripts from the genic regions represent mostly mRNAs. Out of these, 6400 transcript assemblies were mapped to 4744, 18, and 10 unique transcripts from the genic regions of nuclear, chloroplast, and mitochondrial genomes respectively (Table S1). The rest of the 3199 transcript assemblies were mapped to the intergenic regions of all the three organellar genomes. To know the presence of full-length mRNAs within the maize apoplast, the transcript assemblies were analyzed for the percentage of sequence coverage on the mapped transcript. The transcript assemblies originating from the genic regions showing sequence coverage of more than 90% of the mapped transcript were considered as full-length transcripts (Table S2). Thus the maize apoplast was found to contain both full length as well as partial mRNAs representing all the three genomes (nuclear, chloroplast, and mitochondrial). Out of the 9599 transcript assemblies mapped in total on the maize genome while 93% (8929) assemblies were mapped to the nuclear genome, 1.8% (178) and 5.12% (492) assemblies were mapped to the chloroplast and the mitochondrial genomes respectively (Figure 1). Most interestingly except in the case of the chloroplast, both the nuclear as well as mitochondrial transcript assemblies have a significantly high representation of those mapping to the intergenic region. About 94% of the exRNA transcript assemblies mapped to the mitochondrial genome were mapped to the intergenic region (Figure 1). Functional classification of the identified nuclear transcripts revealed only 1194 transcripts to which different Gene Ontology Biological Process (GO_BP) terms can be assigned. Accordingly, among the transcripts, representatives from 12 major biological processes were obtained with confidence (p<0.05) (Table S3). Most of the transcripts were found to be involved in two different biological processes namely ‘cellular processes’ (GO:0009987) and ‘metabolic processes’ (GO:0008152). Besides these, the other most populated categories obtained were ‘biological regulation’, ‘response to stimulus’, ‘localization’, and ‘signalling’ (Figure 2a). Further insights into the possible biological function of this extracellular RNA pool was obtained by carrying out a statistical overrepresentation analysis of the mapped transcripts with respect to the overall composition of the maize transcriptome. Interestingly biological processes representing different stages of photosynthesis are overrepresented by at least 6 folds in the apoplastic RNA pool of maize. Other overrepresented processes, those worth mentioning here include protein repair and response to stimuli including herbicide, nitrogen levels, and high light intensities (Figure 2b). Infection with *U. maydis* led to an overall induction in the apoplastic RNA transcripts. More than 75% of the transcripts identified to undergo differential expression within the apoplast of maize upon infection with *U. maydis* when compared to uninfection condition, exhibited induced expression (Table S4). Among these transcripts, those involved in the ‘ribosome biogenesis pathway’ are enriched to at least 4 folds with respect to the total differentially expressed transcripts upon *U. maydis* infection when compared to the apoplastic RNA pool of uninfected maize as obtained through a statistical overrepresentation analysis (Figure 3a). Among the transcripts categorized under the ‘ribosome biogenesis pathway’ in *U. maydis* infected maize exRNA are included those coding for ribosomal proteins and both 5.8S and 18S rRNAs (Table S5). Figure 3b shows the relative abundance of the exRNA transcripts corresponding to ‘ribosome biogenesis pathway as obtained from the RNA sequencing data. We also tested the relative expression of a set of exRNA transcripts coding for ribosomal proteins in the exRNA pool of maize with *U. maydis* SG200 WT infection and uninfected maize using real time PCR analysis. Figure 3c shows an induced expression of all the tested transcripts as expected from the RNA sequencing data. Since there are no known standard apoplastic transcripts in maize with a constitutive expression we used the expression level of linoleate 13S-lipoxygenase10 RNA (NM_001112510.2) that showed uniform transcript abundance in the exRNA sequence data in all of our samples for the normalization of the real-time data. Our observations till this point raise the most obvious question of whether the maize apoplast contains ribosomes. Although at this stage of the study the presence of extracellular ribosomes in maize cannot be confirmed, pieces of evidence prevail indicating intact ribosomes even tRNAs within the RNase inhibitor-treated cell-conditioned media of human breast cancer cell line, MCF-7 ^7^. This hints toward the possibility of extracellular translation events. However, under our experimental condition and sequencing parameters, no tRNA sequences could be detected within the exRNA transcriptome. However, we did find some of the transcripts coding for tRNA synthetases and ligases (Table S6). Further enrichment of the extracted apoplastic RNA for small RNA species with sizes ranging from 70-150 nucleotides might reveal the presence of tRNAs in maize apoplast as well. Nevertheless, *U. maydis* infection in maize also induced expression of many of the exRNA transcripts involved in mRNA processing including U2 spliceosomal RNA and several other non-coding RNAs (Figure 4, Table S7a). In addition, exRNA transcripts coding for various splicing factors were also detected within maize apoplast (Table S7b). Our data revealed few micro-RNA transcripts within the apoplast of uninfected and infected maize as well. These include miR156d, miR398a, miR11969, and miR11970 (Table S8). Some of these detected miRNAs have proven roles in plant stress response. miR398 for instance is involved in a wide variety of stress responses including oxidative stress, water deficit, salt stress, and biotic stress involving plant pathogenic microbes ^8^. In a recent study natural antisense transcripts of miR398 have been shown to regulate the miR398 mediated thermotolerance in *Arabidopsis* and *Brassica* ^9^. Likewise, miR156 has been demonstrated to play regulatory roles in the process of anthocyanin biosynthesis and flowering in Poplar and Arabidopsis respectively ^10, 11^. miR11969 and miR11970 are involved in the process of meiosis in *Z. mays* ^12^. Although our sequencing data revealed the presence of these important miRNAs within the apoplast of maize no significant difference in their expression was noted upon infection with *U. maydis*. Further analysis of the abundance of these transcripts in small RNA enriched exRNAs from uninfected and *U. maydis* infected maize might provide detailed insights in the future. Although most of the exRNA transcripts detected within the maize apoplast were induced upon *U. maydis* infection about 25 % showed reduced expression as well. Among these are included the ones that code for proteins involved in photosynthesis (Table S9). A group of 38 transcripts all coding for proteins involved in different stages of photosynthesis are at least 3 folds enriched among the differentially expressed transcripts in *U. maydis* infected *Z. mays* when compared to the total apoplastic transcripts obtained in uninfected maize in a statistical overrepresentation analysis (Figure S2). Among the other downregulated processes during *U. maydis* infection are included biological regulation, cellular processes, and response to a stimulus. We have previously demonstrated the nucleolytic activities of a set of two secreted T2 type ribonucleases, Nuc1 and Nuc2 on the exRNA pool of *Z. mays*. We hypothesized and subsequently proved the role of these nucleases in the scavenging of *Z. mays* exRNA as a phosphate source during in planta growth of the pathogen ^6^. However, we could not exclude any involvement of the secretary nucleases in regulating the possible exRNA-mediated cell-cell communication in *Z. mays*. So, in the present study together with uninfected and *U. maydis* SG200 WT infected *Z. mays* apoplastic RNA we also sequenced the exRNA from SG200Δ*nuc1*Δ*nuc2* infected maize. A total of 1655 transcript assemblies showing differential expression between SG200 WT infected and SG200Δ*nuc1*Δ*nuc2* infected maize was obtained that mapped to 1276 unique transcripts in the maize nuclear genome (Table S10). About 50% of these transcripts showed reduced abundance in SG200 WT infected maize apoplast compared to SG200Δ*nuc1*Δ*nuc2*. Considering the fact that *U. maydis* SG200 WT cells possess active Nuc1 and Nuc2 in the apoplast while SG200Δ*nuc1*Δ*nuc2* infected apoplast lack them, the nucleolytic activity within SG200Δ*nuc1*Δ*nuc2* infected maize apoplast is less compared to SG200 WT infected one. The set of transcripts that are found in reduced abundance in SG200 WT infected maize apoplast might therefore be the ones upon which the ribonucleases have acted. Among the transcripts that are abundantly present within the apoplast of maize infected with SG200Δ*nuc1*Δ*nuc2* are included those that code for some of the important signalling kinases including MAP kinase, calcium-dependent protein kinase proteins, and LRR receptor-like serine/threonine protein kinases (Table S11). Kinases belonging to these families have been demonstrated to exhibit key roles in plant development as well as a stress response in many instances. For example, many MAP kinases have been shown to be involved in the defense response of plants to different stresses including pathogen attack, light stress, salt stress, drought stress, and metal stress ^13^. In maize, ZmMPK3, ZmMPK17, ZmMKK4, and ZmMKK1 all belong to the MAP kinase family and have been found to be involved in the response of maize to multiple environmental stresses ^14-17^. In a recent study, a calcium-dependent protein kinase ZmCPK11 from maize was found to be a part of the ‘touch’ and ‘wound-induced defense responsive pathway of maize ^18^. Likewise, many transcripts coding for different phosphatases has also been found to be abundantly expressed in SG200Δ*nuc1*Δ*nuc2* apoplast compared to the SG200 WT infected apoplast. Most interestingly we also obtained transcripts coding for a group of transcription factors including WRKY53 which is a known regulator of plant senescence as well as stress response ^19, 20^. Although, we do not exclude the possibility that the higher abundance of these exRNAs in the SG200Δ*nuc1*Δ*nuc2* infected apoplast is due to plant’s enhanced response against the attenuated pathogen lacking secretary nucleases. This study presents several key transcripts with a potential role in the stress response that is exported out of the cell into the apoplast under both uninfection as well as infection conditions. But whether these transcripts contribute to the growth, development, and stress response in the host plant maize and the mechanism thereof remains elusive. Studies have shown the transport of extracellular mRNAs primarily enclosed within EVs into host cells that are capable of subsequent translation events leading to the synthesis of the encoded proteins ^21^ in animal cells. Although the existence of a similar mechanism in plants needs extensive research, the hypothesis can explain the biological function of the exRNAs in host plants very well. The full-length translation compatible mRNAs might be exported by a stressed plant cell within the apoplast for subsequent translocation to adjacent unstressed cells. These adjacent cells then are primed for defending the upcoming stressed condition through the translation of the received transcripts into the respective proteins. Like for instance a MAP kinase transcript that codes for a stress-responsive MAP kinase may be secreted by a plant cell in response to environmental stress. This particular MAP kinase transcript if taken up by an adjacent cell and translated, then cell would remain prepared to encounter the stress in advance that would eventually spread to this new cell. However, a pathogen secreted nuclease if can target this MAP kinase transcript and efficiently prevent the spread of ‘immunity’ can significantly contribute towards pathogen virulence. However, our transcriptome data also revealed exRNA transcripts that can be treated as signatures of possible extracellular protein translation events in maize. These include transcripts coding for various proteins involved in ribosome biogenesis and functioning including different ribosomal proteins and rRNAs. Moreover, we also obtained exRNA transcripts coding for histidyl and prolyl tRNA synthetases and alanine and threonine tRNA ligases. Consistent with both the hypothesis our data showed an overrepresentation of RNA binding molecular function within the apoplastic transcripts in uninfected maize in relation to the total transcriptome of the host plant. A group of 9 transcripts coding for glycine rice RNA binding proteins were detected in the apoplastic RNA pool of uninfected *Z. mays* (Table S12). Five of these transcripts were found to exhibit increased abundance during *U. maydis* infection. A translation of these templates can build up a pool of proteins protecting the integrity of the exRNAs. In summary, maize apoplast demonstrates one of the host cellular compartments loaded with RNA transcripts indicating potential involvement in host defence against pathogen invasion involving as yet unknown mechanism. It is evident from our study that *U. maydis* infection induces an increased abundance of the majority of the exRNA transcripts in maize. However, how the pathogen copes up in this apoplastic environment loaded with maize transcripts many of which code for defence proteins is yet to be studied. Also, any role of extracellular ribonucleases of *U. maydis* in the modulation of exRNA mediated biological function in maize remains to be revealed.

**Figure 1.**
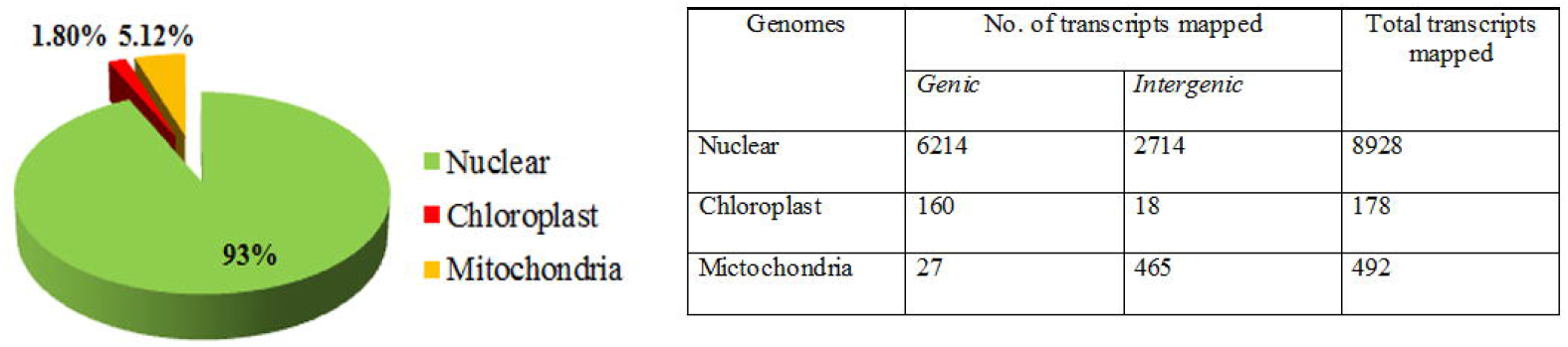
Distribution of exRNA transcript assemblies from uninfected maize apoplastic RNA sequencing data over the nuclear, chloroplast and mitochondrial genomes of maize. Pie chart showing the relative contribution of transcript assembles mapped to the nuclear, choloroplast and mitochondrial genomes of maize. The table lists the number of transcript assemblies mapped to the different genomes.

**Figure 2.**
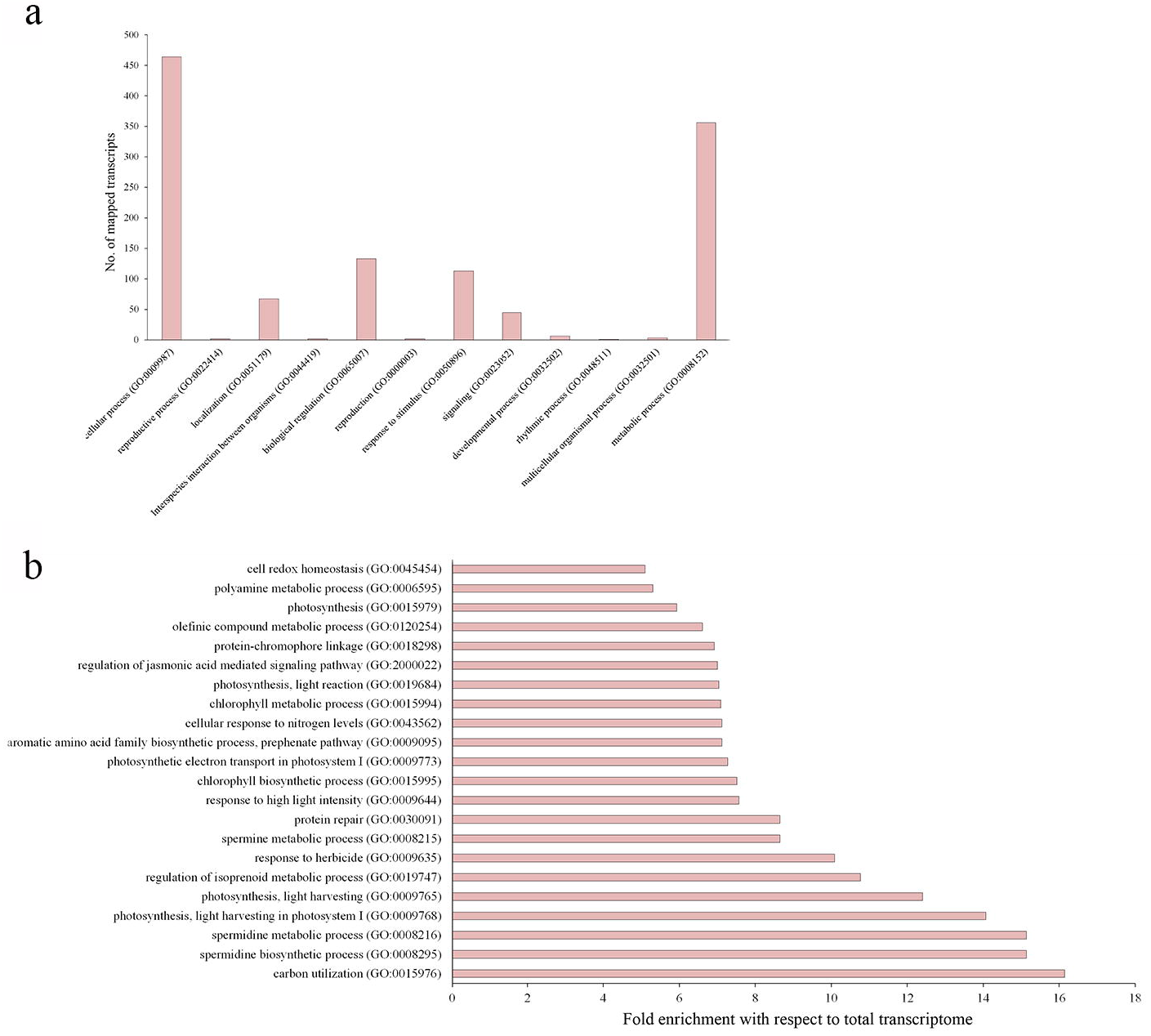
Functional classification of mapped exRNA transcripts from uninfected maize apoplast. (a) Graph showing the number of unique exRNA transcripts mapped to the nuclear genome of maize categorised under indicated Gene Ontology biological processes (GO_BP) categories. (b) Graph showing the overrepresented biological processes in the exRNA transcriptome of uninfected maize with respect to the total transcriptome. Statistical overrepresentation analysis was carried out in Panther database (available at http://www.pantherdb.org/) with the uninfected maize exRNA transcript using maize total transcript as the reference.

**Figure 3.**
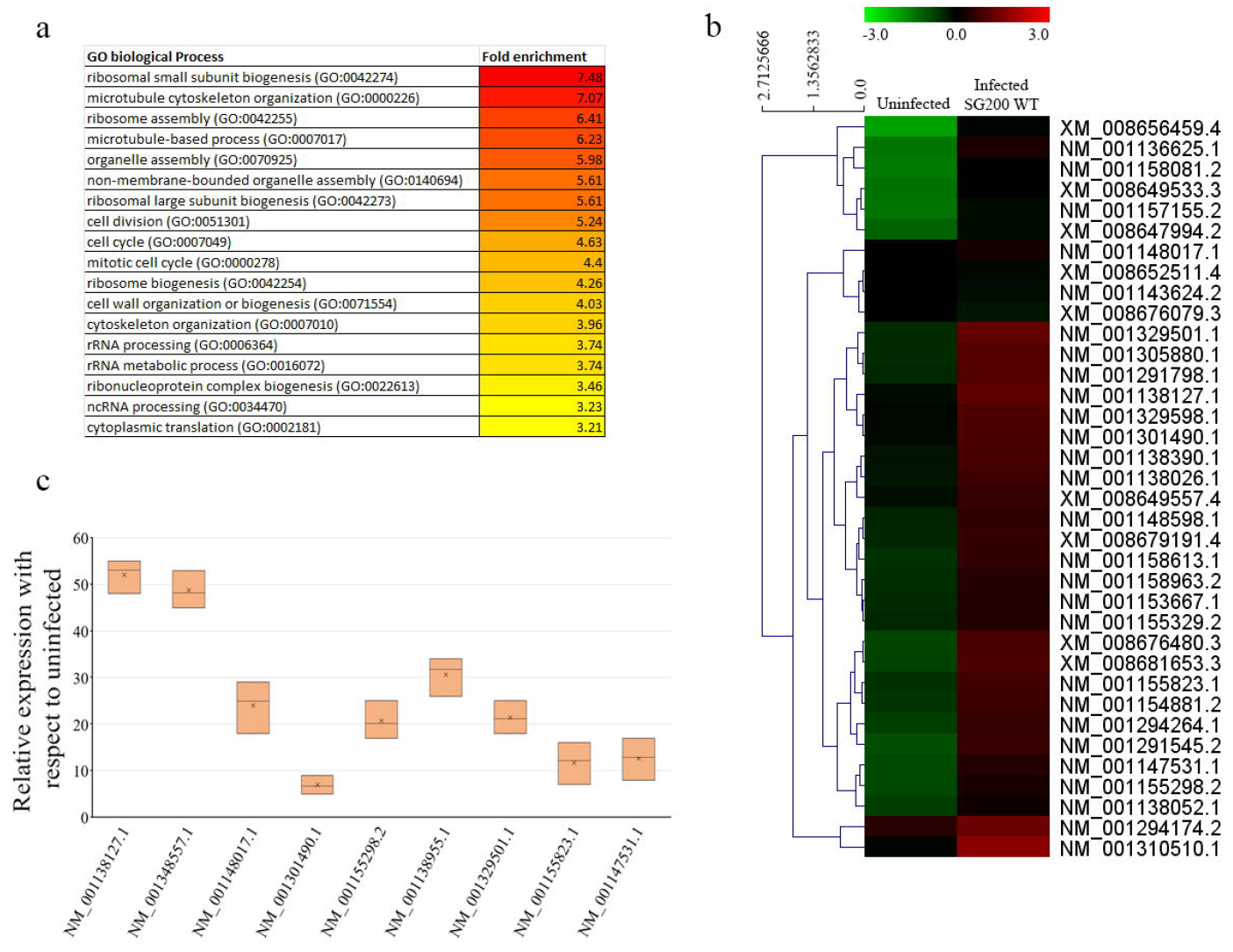
ExRNA transcripts corresponding to ribosome biogenesis pathway are overrepresented in *U. maydis* SG200 WT infected maize apoplast. (a) Graph showing different Gene Ontology biological processes (GO_BP) that are overrepresented within the exRNA transcripts from *U. maydis* infected maize apoplast with respect to uninfected apoplast. The degree of fold enrichment is colour coded. (b) Heat map showing the relative abundance of exRNA transcripts corresponding to the ‘ribosome biogenesis pathway’ in the apoplast of uninfected maize when compared to infection with *U. maydis* SG200 WT. (c) Graph showing the relative expression of each of the selected candidate exRNA transcripts coding for ribosomal proteins in *U. maydis* SG200 WT infected maize apoplast when compared to the expression of the respective transcripts in uninfected apoplast in a quantitative real time PCR experiment. The expression of linoleate 13S-lipoxygenase10 RNA (NM_001112510.2) that showed uniform transcript abundance in the exRNA sequence data in all of the samples was used for the normalisation of the real time data.

**Figure 4.**
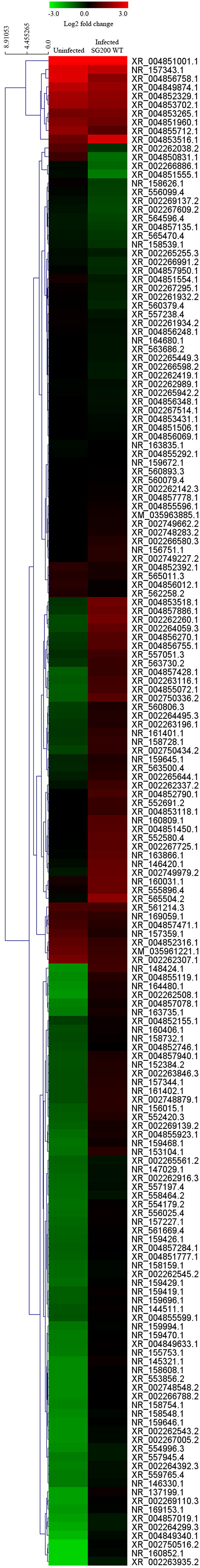
Heat map showing the relative abundances of the non-coding exRNA transcripts within the apoplast of uninfected maize when compared to that within *U. maydis* SG200 WT infected maize.

## Acknowledgement

AG thanks Bose Institute Intramural grant for funding the study. AR and RM received fellowship from DBT-JRF program.

## Author contribution

AG conceived the study. DM, AR, RM and UA performed experiments. NPS and DC analysed the next generation sequencing data. AG carried out all the in-silico analysis related to functional enrichment of the identified exRNA transcripts in different samples. AG wrote the paper with inputs from all the authors.

## Conflict of interest

The authors declare that they have no conflicts of interest.

## Data availability statement

The data that support the findings of this study are available from the corresponding author upon reasonable request. The ExRNA transcriptome data reported in this manuscript were submitted to NCBI SRA database with the accession PRJNA843195.

## Notes

https://www.ncbi.nlm.nih.gov/sra/PRJNA843195

### Competing Interest Statement

The authors have declared no competing interest.

