## supplemental method and figure for "The extracellular RNA pool within *Zea mays* apoplast: composition and differential expression during *Ustilago maydis* infection"

**SUPPLEMENTARY INFORMATION**

- Materials and methods
- Figure S1. Comparative gel profile of leaf apoplastic exRNA and total RNA.
- Figure S2. Statistical overrepresentation analysis of exRNA transcripts showing reduced abundance in *U. maydis* SG200 WT infected Z. mays apoplast compared to uninfected apoplast.
- Table S1. Mapped exRNA transcripts from the apoplast of uninfected Z. mays. (excel file)
- Table S2. Full length apoplastic RNA transcripts from uninfected maize. (excel file)
- Table S3. Biological processes to which the the mapped transcripts from uninfected maize apoplastic RNA pool are assigned. (excel file)
- Table S4. Differentially expressed transcripts in uninfected maize apoplast with respect to infection with Ustilago maydis SG200. (excel file)
- Table S5. Maize apoplastic transcripts induced during U. maydis SG200 infection showing overrepresented biological processes involved in 'ribosome biogenesis pathway' with respect to total apoplastic trancriptome of maize. (excel file)
- Table S6. ExRNA transcripts detected in maize apoplast under infection and uninfection condition coding for proteins involved in tRNA maturation and charging with specific amino acids. (excel file)
- Table S7a. Non coding exRNA transcripts detected in U. maydis infected maize apoplast (excel file)
- Table S7b. ExRNA transcripts coding for splicing factors detected within apoplastic transcriptome of maize under infection and uinfection condition. (excel file)
- Table S8. miRNAs detected within exRNA transcripts from maize apoplast under uninfection and infection conditions. (excel file)
- Table S9. Relative abundance of the exRNA transcripts coding for proteins involved in different stages of photosynthesis in uninfected Z. mays apoplast with respect to U. maydis SG200 WT infected Z. mays apoplast. (excel file)
- Table S10. Relative abundance of the exRNA transcripts in U. maydis SG200 WT infected Z. mays apoplast with respect to U. maydis SG200∆nuc1∆nuc2 infected Z. mays apoplast. (excel file)
- Table S11. ExRNA transcripts with Panther/GO functional classification showing increased abundance in SG200∆nuc1∆nuc2 infected maize apoplast with respect to SG200 WT infected apoplast. (excel file)
- Table S12. ExRNA transcripts with RNA binding molecular function detected in uninfected maize apoplast. (excel file)
- Table S13. Primers used in this study. (excel file)

**Materials and methods**

*Fungal Strains and host plant*

The *Ustilago maydis* solopathogenic strain SG200 [1] and a double deletion mutant of two T2 type secreted ribonucleases Nuc1 and Nuc2 in SG200, SG200Δ*nuc1Δnuc2* [2] are used in this study. All the plant experiments were carried out in B73 variety of maize.

*Infection of maize plants*

Overnight grown cultures of different *U. maydis* strains were inoculated in YEPSL media [2] at an OD_600_ of 0.2 and incubated at 28^0^C at 200 rpm till the OD_600_ reached 0.8. The cell cultures were then centrifuged at 3,000 g for 5 min following which the supernatants were discarded. The resulting cell pellets were resuspended in sterile water to a final OD_600_ value of 1.0. These cell suspensions were used to infect maize seedlings through syringe-infection method as described previously [1].

*Extraction of Apoplastic Fluid*

Apoplastic fluid was extracted from the maize leaf tissue as described previously [2]. Briefly, maize leaves from either 12 days old uninfected seedlings or 5 days post infection (5 dpi) with SG200 WT or SG200∆*nuc1*∆*nuc2* were cut into 7-8 cm long pieces and infiltrated under vacuum with infiltration buffer IB (20 mM MES pH 6.0, 2 mM CaCl_2_, 0.01 M NaCl) at 80 kPa for 4×20 min cycles. After infiltration excess buffer was wiped gently from the leaf surface. The leaves were bundled and packed into 20 ml needleless syringes. The syringes were placed into 50 ml centrifuge tubes and apoplastic fluid was extracted by centrifugation at 2,500 g for 20 min at 4^0^C.

*Total RNA Isolation from the Apoplastic Fluid*

The apoplastic fluid obtained was subjected to centrifugation at 12,000 g for 30 min at 4^0^C for the removal of any fine suspended particles. The supernatant was then mixed with 0.1 volumes of 3 M sodium acetate, pH 5.0 and 0.05 μg/μl RNase free glycogen (Thermo scientific). To the mixture 1 volume of ice-cold isopropanol was added, vortexed briefly and incubated for 60 min at -20^0^C for RNA precipitation. Following incubation, the mixture was centrifuged at 12,000 ×g for 30 min at 4^0^C. The RNA pellet thus obtained was resuspended in 1 ml of TRIzol reagent (Ambion) for further purification of the extracted RNA following manufacturer’s protocol. Total RNA from the apoplastic fluid thus obtained was incubated with DNase I (NEB) at 37^0^C for 30 min and was subjected to a second step purification using TRIzol reagent. The RNA pellet thus obtained was washed in 75% ethanol (V/V) in DEPC water and finally resuspended in 1X gel loading dye and run on a 8% urea PAGE as described in [2]. For real time expression analysis the extracted RNA pellets were stored in 75% ethanol (V/V) in DEPC water at -80^0^C for future use.

*Next generation sequencing of the apoplastic RNA samples*

Apoplastic RNA extracted from either uninfected or *U. maydis* infected maize were sequenced using Illumina HiSeqX10 sequencing platform. TruSeq stranded mRNA library preparation protocol was followed to prepare the sequencing libraries. The raw sequencing data were submitted to NCBI SRA database with the accession (PRJNA843195).

*Analysis of RNA sequencing data*

Raw Illumina reads sequenced from the exRNA samples were quality checked with FastQC, adapter sequences and low quality regions (Q30) were trimmed at the end with NGSQC Toolkit (v2.3.3) [3]. Further, the quality reads were mapped to the maize reference genome (Zm-B73-REFERENCE-NAM-5.0), chloroplast genome (X86563.2), and mitochondrial genome (AY506529.1) employing bwa (v0.7.17) with the default parameters [4]. Subsequently, the reads that were mapped in each of the reference is used for their respective trancriptome assembly with Trinity [5]. To check whether the assembled transcripts are coded by a gene or originated from the noncoding intergenic parts, nuclear, chloroplast and mitochondrial genomes were mapped with transcripts from their corresponding transcriptomes with BLAST (v2.12.0+). If the genomic coordinate where the transcript aligns harbour a gene, such transcripts are designated as genic or else intergenic. Those transcript which show >= 90% sequence similarity with maize reference mRNA sequences were considered complete. RSEM [6] was used to estimate the FPKM values based on bowtie2 [7] alignment of reads to the transcriptome and the differential expression analysis was performed by the DESeq2 package [8]. With a count-based approach, this R package employs the negative binomial distribution to account for both biological and technical variability among the samples. Finally, differentially expressed genes (DEGs) were selected by applying a cut-off of p-value ≤ 0.05, FDR ≤ 0.05 and log_2_ fold change ≥ +1.0 and ≤−1.0. Noncoding RNAs such as miRNA and tRNA among the transcriptome were identified. miRNA sequences obtained from mirbase database (<https://www.mirbase.org/>) [9] was used to map against the transcriptome with bowtie using default parameters and putative tRNAs were predicted with tRNAscan-SE [10].

Quantitative real time PCR

For assessing the relative expression of different identified apoplastic transcripts in infected and uninfected maize, the exRNA isolated from the respective apoplasts were first treated with DNaseI. Following this the DNaseI treated exRNAs were reverse transcribed using Revert Aid reverse transcriptase (Thermo Scientific) and gene specific reverse primers following manufacturer’s protocol. 1.5 µg of exRNA was used as template in each of the cDNA synthesis reactions. Quantitative real time PCR was performed using TB Green Premix Ex Taq (Tli RNaseH Plus) qPCR kit (Takara Biosciences) in a CFX96 real time PCR detection system (BioRad). Atleast three biological replicates were performed for each of the reactions and 2^-∆∆Ct^ method [11] was used to calculate the fold change of the tested genes between different samples. Since there are no known standard apoplastic transcripts in maize with constitutive expression we used the expression level of linoleate 13S-lipoxygenase10 RNA (NM_001112510.2) that showed uniform transcript abundance in the RNA sequence data in all of our samples for the normalisation of the real time data. Sequences of the primers used are listed in Table S13.

*Functional enrichment analysis of the identified and differentially expressed exRNA transcripts*

All the analysis related to functional classification, enrichment and statistical overrepresentation of the identified and differentially expressed exRNA transcripts in maize were carried out in Panther database (available at pantherdb.org) based on the Gene Ontology biological process (GO_BP) terms that can be assigned to each of the mapped transcripts. For statistical overrepresentation analysis of apoplastic RNA from uninfected maize the set of total transcribed genes in maize was used as the reference. However, for the statistical overrepresentation of the differentially regulated exRNA transcripts in maize infected with either *U. maydis* SG200 WT or SG200∆*nuc1*∆*nuc2* the set of apoplastic RNA transcripts from uninfected maize was used as the reference.

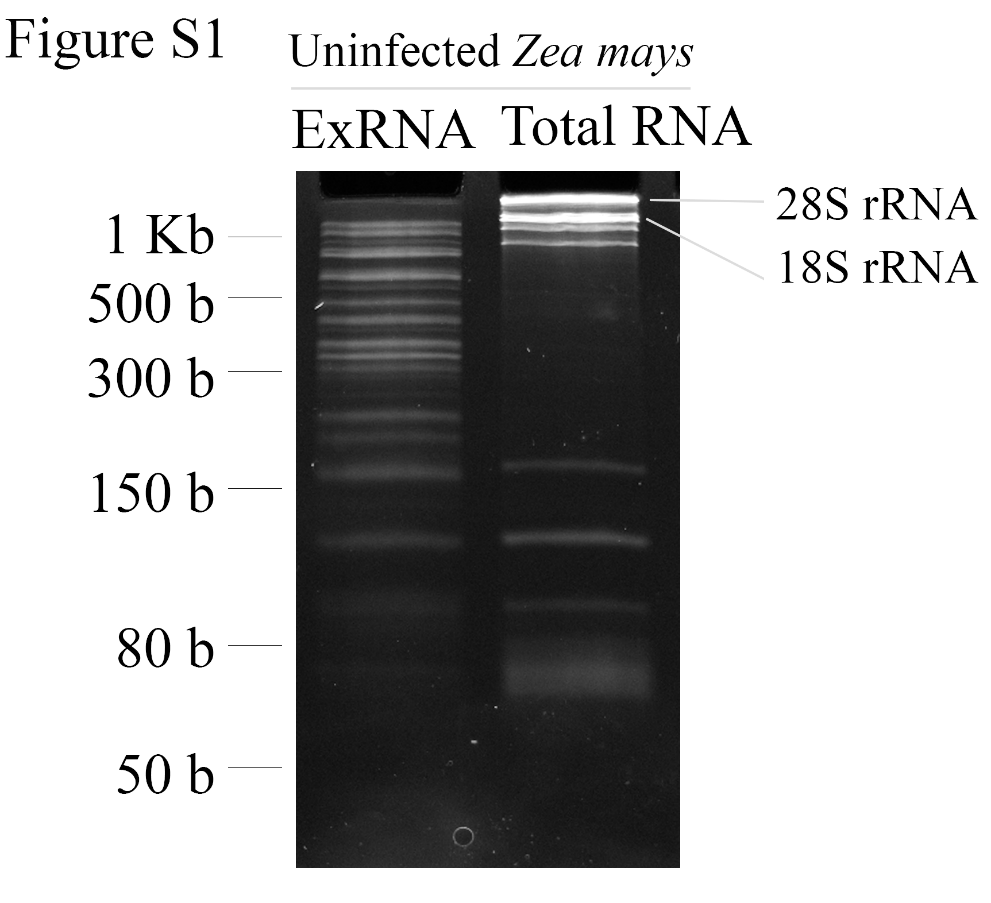

Figure S1. Comparative gel profile of leaf apoplastic exRNA and total RNA. 8% urea polyacrylamide gel showing the profile of the exRNA isolated from the apoplast of maize leaves under normal uninfection condition and the RNA extracted from leaf tissue as a whole.

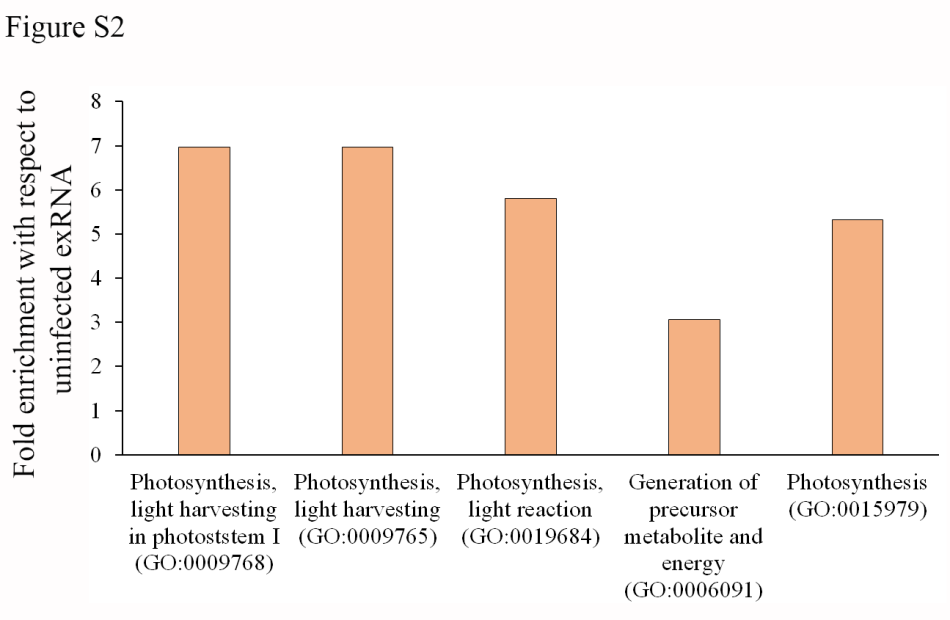

Figure S2. Statistical overrepresentation analysis of exRNA transcripts showing reduced abundance in *U. maydis* SG200 WT infected Z. mays apoplast compared to uninfected apoplast. The total exRNA transcriptome of uninfected *Z. mays* is used as the reference for the analysis.
